# Gene model for the ortholog of *Pi3K21B* in *Drosophila eugracilis*

**DOI:** 10.1101/2025.07.21.665947

**Authors:** Bethany C. Lieser, Ryan M. Bonthius, Clairine I.S. Larsen, Jeffrey S. Thompson, Stephanie Toering Peters

## Abstract

Gene model for the ortholog of *Phosphatidylinositol 3-kinase 21B* (*Pi3K21B*) in the *D. eugracilis* May 2021 (Stanford ASM1815383v1/DeugRefSeq2) Genome Assembly (GenBank Accession: GCF_018153835.1) of *Drosophila eugracilis*. This ortholog was characterized as part of a developing dataset to study the evolution of the Insulin/insulin-like growth factor signaling pathway (IIS) across the genus *Drosophila* using the Genomics Education Partnership gene annotation protocol for Course-based Undergraduate Research Experiences.

---

> *This article reports a predicted gene model generated by undergraduate work using a structured gene model annotation protocol defined by the Genomics Education Partnership (GEP; thegep.org) for Course-based Undergraduate Research Experience (CURE). The following information may be repeated in other articles submitted by participants using the same GEP CURE protocol for annotating Drosophila species orthologs of Drosophila melanogaster genes in the insulin signaling pathway*.

“In this GEP CURE protocol students use web-based tools to manually annotate genes in non-model *Drosophila* species based on orthology to genes in the well-annotated model organism fruitfly *Drosophila melanogaster*. The GEP uses web-based tools to allow undergraduates to participate in course-based research by generating manual annotations of genes in non-model species (Rele et al., 2023). Computational-based gene predictions in any organism are often improved by careful manual annotation and curation, allowing for more accurate analyses of gene and genome evolution (Mudge and Harrow 2016; Tello-Ruiz et al., 2019). These models of orthologous genes across species, such as the one presented here, then provide a reliable basis for further evolutionary genomic analyses when made available to the scientific community.” (Myers et al., 2024).

“The particular gene ortholog described here was characterized as part of a developing dataset to study the evolution of the Insulin/insulin-like growth factor signaling pathway (IIS) across the genus *Drosophila*. The Insulin/insulin-like growth factor signaling pathway (IIS) is a highly conserved signaling pathway in animals and is central to mediating organismal responses to nutrients (Hietakangas and Cohen 2009; Grewal 2009).” (Myers et al., 2024).

“The gene product of the *Pi3K21B* gene (FBgn0020622), p60, was identified through affinity purification based on binding to a phosphorylated peptide in a heterodimer with p110, the *Pi3K92E* gene product (Weinkove et al., 1997). Additional experiments indicated that the p60•p110 complex was present in all life cycle stages in *Drosophila*, and demonstrated protein and lipid phosphatase activity of the complex in vitro (Weinkove et al., 1997). Null alleles were generated in *Drosophila* through P-element mobilization, and homozygous null animals display reduced larval growth with death in the third instar or early pupal stage of development (Weinkove et al., 1999). Further work indicated that the p60 adaptor protein was required for PI3K activity in the insulin signaling pathway, and was involved in determining both cellular and organismal size and growth (Britton et al., 2002; Oldham et al., 2002).” (Backlund et al., 2025).

We propose a gene model for the *D. eugracilis* ortholog of the *D. melanogaster Phosphatidylinositol 3-kinase 21B* (*Pi3K21B*) gene. The genomic region of the ortholog corresponds to the uncharacterized protein XP_017075761.2 (Locus ID LOC108110971) in the D. eugracilis May2021(Stanford ASM1815383v1/DeugRefSeq2) Genome Assembly of *D. eugracilis* (GCF_018153835.1; Kim et al., 2022). This model is based on RNA-Seq data from *D. eugracilis* (SRP008375, PRJNA73487; Chen et al., 2014; Kim et al., 2015; Wang et al., 2012) and *Pi3K21B* in *D. melanogaster* using FlyBase release FB2023_03 (GCA_000001215.4; Larkin et al., 2021; Gramates et al., 2022, Jenkins et al., 2022). “*D. eugracilis* is part of the *melanogaste*r species group within the subgenus *Sophophora* of the genus *Drosophila* (NCBI:txid29029; Pélandakis et al., 1993). It was first described as *Tanygastrella gracilis* by Duda (1924) and revised to *Drosophila eugracilis* by Bock and Wheeler (1972). *D. eugracilis* is found in humid tropical and subtropical forests across southeast Asia (https://www.taxodros.uzh.ch, accessed 1 Feb 2023).” (Morgan et al., 2022).

## Synteny

The reference gene, *Pi3K21B*, occurs on chromosome 2L in *D. melanogaster* and is flanked upstream by *U2 small nuclear riboprotein auxiliary factor 38* (*U2af38*) and *Stress induced phosphoprotein 1* (*Stip1*) and downstream by *Phospholipase C at 21C* (*Plc21C*) which nests *CG11912, CG11911, CG33127, CG43755* and *CG31921*, and further downstream by *Equilibrative nucleoside transporter 1* (*Ent1*). The *tblastn* search of *D. melanogaster* Pi3K21B-PE (query) against the *D. eugracilis* (GenBank Accession: GCF_018153835.1) Genome Assembly (database) placed the putative ortholog of *Pi3K21B* within scaffold NW_024573036 (NW_024573036.1) at locus LOC108110971 (XP_017075761.2)— with an E-value of 8e-137 and a percent identity of 76.22%. Furthermore, the putative ortholog is flanked upstream by LOC108110743 (XP_017075436.1) and LOC108110741 (XP_017075435.1), which correspond to *U2af38* and *Stip1* in *D. melanogaster* (E-value: 0.0 and 0.0; identity: 98.46% and 93.27%, respectively, as determined by *blastp*; Figure 1A; Altschul et al., 1990). The putative ortholog of *Pi3K21B* is flanked downstream by LOC108110680 (XP_017075294.2), which nests LOC108110687 (XP_017075303.2), LOC108110686 (XP_017075302.2), LOC108110685 (XP_017075301.2), LOC108110683 (XP_041674059.1) and LOC108110681 (XP_017075298.2), and further downstream by LOC108110682 (XP_017075299.1), which correspond to *Plc21C, CG11912, CG11911, CG33127, CG43755, CG31921* and *Ent1* in *D. melanogaster* (E-value: 0.0, 1e-159, 0.0, 1e-170, 2e-172, 0.0 and 0.0; identity: 96.36%, 78.60%, 85.92%, 85.61%, 63.97%, 53.21% and 88.96%, respectively, as determined by *blastp*). The putative ortholog assignment for *Pi3K21B* in *D. eugracilis* is supported by the following evidence: The genes surrounding the *Pi3K21B* ortholog are orthologous to the genes at the same locus in *D. melanogaster* and local synteny is completely conserved, supported by E-values and percent identities, so we conclude that LOC108110971 is the correct ortholog of *Pi3K21B* in *D. eugracilis* (Figure 1A).

**Figure 1.**
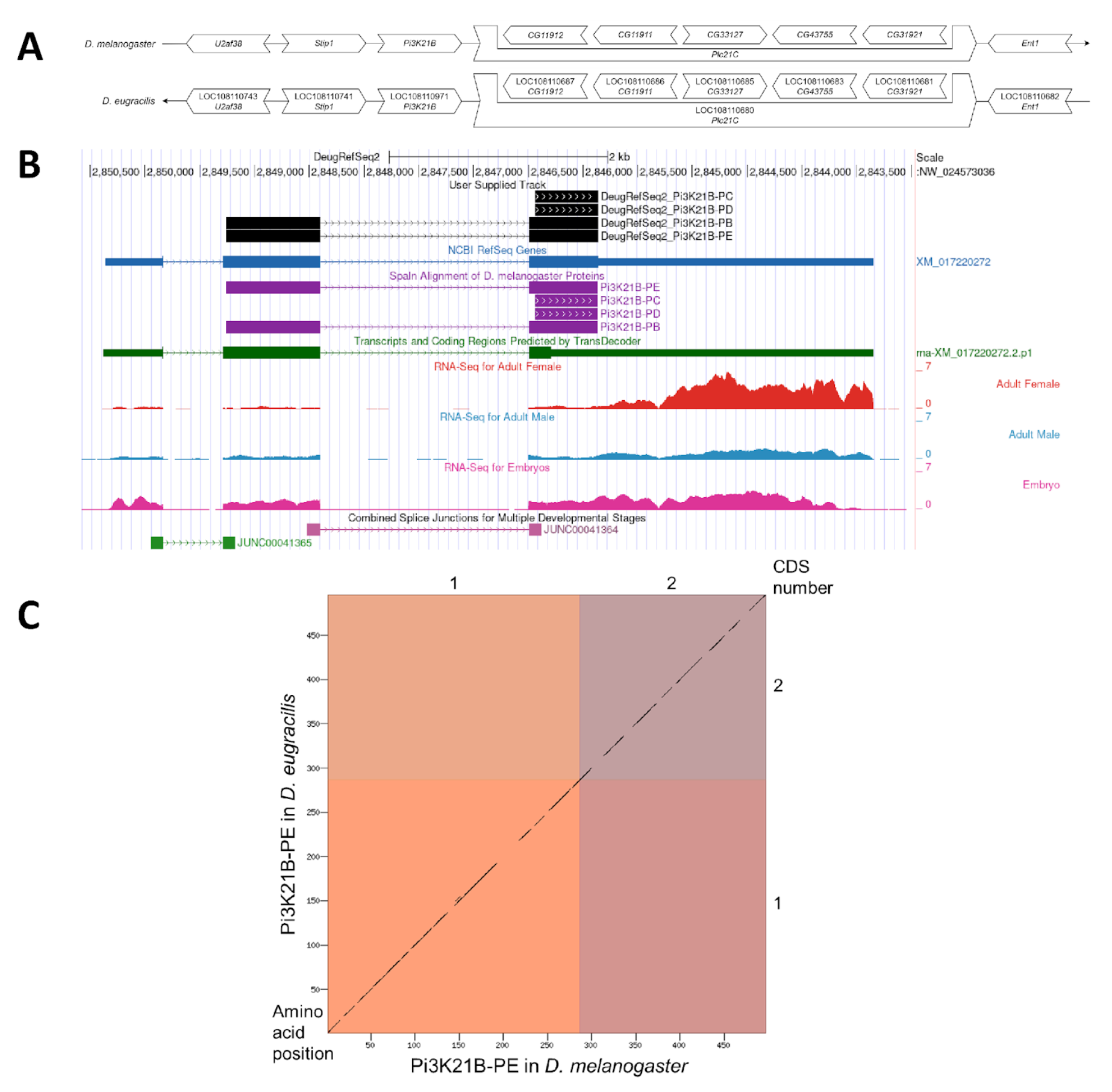
*Pi3K21B* gene model comparison between *Drosophila eugracilis* and *Drosophila melanogaster*. (A) Synteny comparison of the genomic neighborhoods for *Pi3K21B* in *D. melanogaster* and *D. eugracilis*. Thin underlying arrows indicate the DNA strand within which the target gene– *Pi3K21B*–is located in *D. melanogaster* (top) and *D. eugracilis* (bottom). The thin arrow pointing to the right indicates that *Pi3K21B* is on the positive (+) strand in *D. melanogaster*, and the thin arrow pointing to the left indicates that *Pi3K21B* is on the negative (-) strand in *D. eugracilis*. The wide gene arrows pointing in the same direction as *Pi3K21B* are on the same strand relative to the thin underlying arrows, while wide gene arrows pointing in the opposite direction of *Pi3K21B* are on the opposite strand relative to the thin underlying arrows. White gene arrows in *D. eugracilis* indicate orthology to the corresponding gene in *D. melanogaster*. Gene symbols given in the *D. eugracilis* gene arrows indicate the orthologous gene in *D. melanogaster*, while the locus identifiers are specific to *D. eugracilis*. **(B) Gene Model in GEP UCSC Track Data Hub** (Raney et al., 2014). The coding-regions of *Pi3K21B* in *D. eugracilis* are displayed in the User Supplied Track (black); coding CDSs are depicted by thick rectangles and introns by thin lines with arrows indicating the direction of transcription. Subsequent evidence tracks include BLAT Alignments of NCBI RefSeq Genes (dark blue, alignment of Ref-Seq genes for *D. eugracilis*), Spaln of *D. melanogaster* Proteins (purple, alignment of Ref-Seq proteins from *D. melanogaster*), Transcripts and Coding Regions Predicted by TransDecoder (dark green), RNASeq from Adult Females, Adult Males, and Mixed Embryos (red, light blue, and dark pink, respectively; alignment of Illumina RNA-Seq reads from *D. eugracilis*), and Splice Junctions Predicted by regtools using *D. eugracilis* RNA-Seq (SRP008375, PRJNA73487). Splice junctions shown have a minimum read-depth of 10 with 50-99 and 100-499 supporting reads shown in green and pink, respectively. **(C) Dot Plot of Pi3K21B-PE in *D. melanogaster* (*x*-axis) vs. the orthologous peptide in *D. eugracilis* (*y*-axis)**. Amino acid number is indicated along the left and bottom; CDS number is indicated along the top and right, and CDSs are also highlighted with alternating colors. Line breaks in the dot plot indicate mismatching amino acids at the specified location between species.

### Protein Model

*Pi3K21B* in *D. eugracilis* has three CDSs within the genome sequence. The first unique protein sequence (Pi3K21B-PE) is translated from two mRNA isoforms that differ in their UTRs (*Pi3K21B-RE, Pi3K21B-RB*; Figure 1B). The second unique protein sequence (Pi3K21B-PC) is translated from two mRNA isoforms that differ in their UTRs (*Pi3K21B-RC, Pi3K21B-RD*; Figure 1B). Relative to the ortholog in *D. melanogaster*, the CDS number and protein isoform count are conserved. The sequence of Pi3K21B-PE in *D. eugracilis* has 86.29% identity (E-value: 0.0) with the protein isoform Pi3K21B-PE in *D. melanogaster*, as determined by *blastp* (Figure 1C). For the validation of this model, a Variant Call Format (VCF) file was generated to correct a putative error in the DeugRefSeq2 assembly (GCF_018153835.1), supported by high quality RNA-Seq data, of a one nucleotide insertion at position 2846330 on the NW_024573036 scaffold. Coordinates of these curated gene models (Pi3K21B-PE, Pi3K21B-PB, Pi3K21B-PD, and Pi3K21B-PC) are stored by NCBI at GenBank/BankIt (accessions BK068413, BK068414, BK068415, and BK068416, respectively). This gene model can be viewed in the *D. eugracilis* genome at this <u>TrackHub</u>.

## Methods

“Detailed methods including algorithms, database versions, and citations for the complete annotation process can be found in Rele et al. (2023). Briefly, students use the GEP instance of the UCSC Genome Browser v.435 (https://gander.wustl.edu; Kent WJ et al., 2002; Navarro Gonzalez et al., 2021) to examine the genomic neighborhood of their reference IIS gene in the *D. melanogaster* genome assembly (Aug. 2014; BDGP Release 6 + ISO1 MT/dm6). Students then retrieve the protein sequence for the *D. melanogaster* target gene for a given isoform and run it using *tblastn* against their target Drosophila species genome assembly (*D. eugracilis* (GCF_018153835.1; Kim et al., 2022)) on the NCBI BLAST server (https://blast.ncbi.nlm.nih.gov/Blast.cgi, Altschul et al., 1990) to identify potential orthologs. To validate the potential ortholog, students compare the local genomic neighborhood of their potential ortholog with the genomic neighborhood of their reference gene in *D. melanogaster*. This local synteny analysis includes at minimum the two upstream and downstream genes relative to their putative ortholog. They also explore other sets of genomic evidence using multiple alignment tracks in the Genome Browser, including BLAT alignments of RefSeq Genes, Spaln alignment of D. melanogaster proteins, multiple gene prediction tracks (e.g., GeMoMa, Geneid, Augustus), and modENCODE RNA-Seq from the target species. Genomic structure information (e.g., CDSs, intron-exon number and boundaries, number of isoforms) for the *D. melanogaster* reference gene is retrieved through the Gene Record Finder (https://gander.wustl.edu/∼wilson/dmelgenerecord/index.html; Rele et al., 2023). Approximate splice sites within the target gene are determined using *tblastn* using the CDSs from the *D. melanogaste*r reference gene. Coordinates of CDSs are then refined by examining aligned modENCODE RNA-Seq data, and by applying paradigms of molecular biology such as identifying canonical splice site sequences and ensuring the maintenance of an open reading frame across hypothesized splice sites. Students then confirm the biological validity of their target gene model using the Gene Model Checker (https://gander.wustl.edu/∼wilson/dmelgenerecord/index.html; Rele et al., 2023), which compares the structure and translated sequence from their hypothesized target gene model against the *D. melanogaster* reference gene model. At least two independent models for this gene were generated by students under mentorship of their faculty course instructors. These models were then reconciled by a third independent researcher mentored by the project leaders to produce the final model presented here. Note: comparison of 5’ and 3’ UTR sequence information is not included in this GEP CURE protocol.” (Gruys et al., 2025)

## Supporting information

Gene model data files

## Extended Data

File Description: “Zip file containing FASTA, PEP, and GFF” Resource Type: “Model”

## Acknowledgements

We would like to thank Wilson Leung for developing and maintaining the technological infrastructure that was used to create this gene model and Laura K. Reed for overseeing the project. Thank you to FlyBase for providing the definitive database for *Drosophila melanogaster* gene models. FlyBase is supported by grants: NHGRI U41HG000739 and U24HG010859, UK Medical Research Council MR/W024233/1, NSF 2035515 and 2039324, BBSRC BB/T014008/1, and Wellcome Trust PLM13398.

